# Chemical characterization, biological assessment and molecular docking studies of essential oil of *Ocimum viride* for potential antimicrobial and anticancer activities

**DOI:** 10.1101/390906

**Authors:** Madhulika Bhagat, Monica Sangral, Khushboo Arya, Rafiq A. Rather

**Affiliations:** School of Biotechnology, University of Jammu, Jammu, India

**Keywords:** Ocimum viride, essential oil (EO), thymol, antibacterial, molecular docking, apoptosis

## Abstract

*Ocimum viride* (family: Lamiaceae) is a medicinally important aromatic plant that grows widely in north western Himalayan range of Indian subcontinent. Essentials oils (EOs) and purified aromatic compounds derived from plants of genus *Ocimum* have long been used in traditional system of medicine to treat various chronic disorders. In this study we made an attempt to assess the chemical composition of essential oil (EO) obtained from *Ocimum viride* for potential antimicrobial and anticancer properties. Gas chromatography-mass spectrometry (GCMS) analysis revealed that EOs of aerial parts (leaves) of *Ocimum viride* contain high amounts of oxygenated monoterpenes, thymol and gamma terpinene. Notably, thymol (~50%) and γ-terpinene (~18%) were identified as the most abundant components of the oil. EOs showed most prominent antibacterial effect against *Bacillus subtilis* and *in silico* molecular docking analyses of antibacterial action against bacterial cell wall of *Bacillus subtilis* showed interaction of thymol with Sec A protein of *Bacillus subtilis* (binding energy of **-**15 kcal/mol) with active site Lys284, Trp275, Leu269, Arg19, Glu277, pro270. While, *in vitro* cytotoxic effect of EO against six human cancer cell lines showed maximum effect with IC_50_ value of ~0.034 ± 0.001μL/ mL against HT-29 colon cancer cell line. DNA fragmentation analysis and cell cycle analysis revealed that EO inhibits the growth of HT-29 colon cancer cells probably through induction of unrepairable DNA damage and subsequent cell death. Taken together, our results indicate that EO possesses potent antimicrobial and anticancer properties, and may find applications in bacterial growth inhibition and cancer therapeutics.

## Introduction

Essential oils are rich sources of complex mixtures of biologically active substances and medicinally important bioactive organic compounds. They have been used for many purposes such as in cosmetics, pharmaceuticals, perfumes, food products, alcoholic beverages, insecticides, antiseptics etc. They are composed of volatile secondary metabolites having excellent spectrum of biological activities such as antimicrobial, analgesic, anti-inflammatory, antioxidant, anticancer properties etc [1-6]. Furthermore, their pleasant aroma, relatively low toxicity and complex mixtures of biologically active substances offer potential novel template molecules for the treatment of various diseases [7].

The genus *Ocimum* (Family: Lamiaceae formerly Labiatae) has been placed among the most important aromatic plants with medicinal properties and recorded as an extreme source of many naturally occurring essential oils and aromatic compounds. The genus *Ocimum* L. belonging to family Lamiaceae (sixth largest family), collectively called as basil is an important economic and medicinal herb, widely distributed in tropical, subtropical, and warm temperate regions of the world known to contains between 50 to 150 species of herbs and shrubs [8, 9]. In India, nine of these reported species have been identified so far mainly confined to tropical and peninsular regions, recorded with numerous medicinal properties cardiopathy, homeopathy, asthma, bronchitis, vomiting, skin diseases etc [10, 11].

*Ocimum viride* Willd. (Van Tulsi), is an exotic West African species. The plant is perennial, erect, much branched, under shrub with elliptic lanceolate brownish green leaves. Flowers are pale yellow. Seeds are globose, brownish and non-mucilaginous. Traditionaly, plant is extensively used as a poultice for rheumatism and lumbago. A decoction of leaves is used in fever and cough. The fresh juice of leaves is used for catarrh and as eye drops for conjunctivitis [12]. Essential oil is pale yellow, viscous with characteristic odour of Thymol, having pungent and spicy flavour. It is extensively used in perfume, flavour and pharmaceutical products and as a powerful antiseptic, anti-oxidant, preservative and disinfectant. It is also used in compounding of synthetic essential oil besides as a starting material for making synthetic menthol and biological activities are reported like antioxidant, antibacterial, antifungal, free radical scavenging, anticancer activities [12, 13, 14, 15, 16, 17]. Although, some information regarding the activity are available but mechanisms of action of essential oil of *Ocimum viride* underlying are still limited. Therefore, the purpose of this study is to isolate the essential oil, elucidate the chemical profile of aerial part of *Ocimum viride* as well as to establish its potential for *in vitro* antibacterial and cytotoxic properties for exploring its potential of pharmacological activity in food and medicine.

## Material and methods

### Collection and isolation of essential oil

Plant material was collected from the farm of CSIR-Indian Institute of Integrative Medicine, Jammu, India. The aerial parts (leaves) of the *Ocimum viride* were harvested at inflorescence initiation stage. EO was extracted immediately after collection of plant material by hydro-distillation using modified clevenger apparatus for 2 h 30 min [18]. The oil samples were dried over anhydrous sodium sulphate (Na_2_SO_4_) and stored at 4 ͦ C for future analysis.

### GC-MS analysis

Analysis of the essential oil was carried out at CSIR-Indian Institute of Integrative Medicine, India, using GC-MS 4000 (Varian, USA) system equipped with CP-SIL 8CB column (30 m×0.32mm i.d., 1μm film thickness). Injector temperature was maintained at 230°C. Oven temperature program used was holding at 60 °C for 5 min, heating to 250 °C at 3°C/min and keeping the temperature constant at 250 °C for 10 min. Helium was used as a carrier gas at a constant flow of 1.0 ml/min and an injection volume of 0.20 μL was employed. The MS scan parameters included electron impact ionization voltage of 70eV, a mass range of 40-500 m/z. The identification of components of the essential oil was based on comparison of their mass spectra with those of NIST05 (version 2.0) library.

### Antibacterial activity assay

The antibacterial activity of leaf essential oil was determined by agar well diffusion method [19]. The bacterial isolates were first grown by streaking on a nutrient agar plate and incubated for 24 hours at 37 °C before use. Cell suspension was prepared by dissolving loop full of culture from the culture plate in autoclaved nutrient broth. Four wells were then bored into each agar plate using a sterile 5 mm diameter cork borer and filled with 20μl solution of 1μL/mL essential oil. Plates were incubated at 37 °C for 24 hours and later observed for zones of inhibition. The effect of EO was compared with that of positive control chloramphenicol to determine the sensitivity of bacterial growth. Each sample was used in duplicate for determination of antibacterial activity.

### Molecular docking studies

Protein structures were retrieved from RCSB PDB in .pdb text format and various possible binding sites for thymol were predicted from COACH server. PyMOL, a molecular visualization tool, was used to identify desired protein sequences in the target protein for binding affinities with thymol. Different analogs of thymol were also searched in ZINC database and analysed for their potential to bind Sec A protein. The structure of thymol was retrieved from the Zinc Database in .mol2 format. The drug-like property of thymol was measured using Lipinski filter. The protein and ligand molecule were imported in Molegro Virtual Docker and visualized for drug interaction sites by using Molegro visualise.

### Cytotoxicity assay

Cytotoxic effect of EO against human cancer cell lines was measured by tetrazolium-based colorimetric assay which measures the reduction of the tetrazolium salt MTT (3-[4,5-dimethylthiazol-2-yl]-2,5-diphenyltetrazolium bromide) into a blue formazan product, mainly by the activity of the mitochondrial cytochrome oxidase and succinate dehydrogenase [20]. 100 μL of cells suspension were plated at a density of approximate 2 ×10^4^ cells per well in a 96-well plate, and were maintained at 37°C in a 5% CO_2_ humid incubator for 24 hours. Different concentrations of EO were added to each group (triplicate wells) and were incubated for 24h, followed by addition of l0 μL (5mg/mL) of MTT dye solution to each well for 4 hours. After removal of the unused MTT dye, cells were treated with 100 μL DMSO and the absorbance at 570 nm was measured using ELISA reader. The cytotoxicity was calculated relative to the control (treated with 0.1% DMSO) and expressed as the concentration of drug inhibiting cell growth by 50% (IC_50_). All tests were run in triplicates.

### DNA fragmentation analysis

HT-29 colon cells after treatment with different concentrations of essential oil (0.1 μL/mL) for 24h were centrifuged at 1500 rpm for 10 min and washed with Dulbecco's PBS. The resultant cell pellet was suspended in 250 μL of lysis buffer (10mM EDTA, 50mM Tris-HCl, 0.5% SDS). Lysed cells were then supplemented with proteinase-K (500μg/mL) at 55°C for 1h and followed by incubation with 200μg/mL DNAase-free RNase at 37 °C for 90 min. The DNA was extracted with 250 μL of phenol:chloroform:isoamyl alcohol (25:24:1) for 1 min and centrifuged at 12,000 rpm for 5 min. The aqueous phase was further extracted with chloroform: isoamyl alcohol (24:1) and centrifuged. DNA was precipitated from aqueous phase with 0.1 volume of 2M NaCl and 2.5 volumes of chilled ethanol and kept at 20 °C overnight. The precipitated DNA was centrifuged at 12,000 rpm for 10 min and dissolved in Tris-EDTA buffer (pH 8.0) and electrophoresed in 1.5% agarose gel at 50V for 90 min. The gel was photographed using Bio-Rad Gel documentation system [21].

### DNA cell cycle analysis

Cell cycle analysis was performed in accordance with the method reported earlier [22]. Colon HT-29 cells (10^6^ cells/ml) in their exponential growing phase were dispended in a six well plate and maintained for 24 h in incubator. Cells were then treated with EO (0.1 μL/mL) and incubated for further 24 h. Cells were trypsnised, centrifuged and washed with PBS. Cells were fixed in 70% ethanol, washed with PBS and then incubated with propium iodide (PI; 25 μg/mL). RNA was removed using RNAase at 37°C for 30 min. The percentages of cells having the sub-G1 population were measured using BD-LSR flow cytometry equipped with blue (488 nm) excitation from argon laser.

### Nuclear morphology analysis

Morphological changes of apoptotic cells were also examined using fluorescence microscopy. The HT-29 cells were treated with the 0.1μg/ml of the essential oil and further cells were harvested, fixed with absolute ethanol, and stained with Hoechst 33258 for 15 min at 37 °C. The cells were then visualized using fluorescence microscopy (Olympus 1×70, Tokyo, Japan) with UV excitation at 300-500 nm. Cells containing condensed and/or fragmented nuclei were considered to be apoptotic cells [23].

## Results

### Chemical composition of essential oil

The light yellow essential oil was obtained by hydrodistillation of *Ocimum viride* aerial part with a yield of 0.81% (v/w). The chemical composition was analysed by GC-MS and result obtained are indicated in chromatogram as Figure 1 **[1].** Majority of the detected compounds were terpenes in nature **(Fig 1)** among others, thymol (~50%), gamma-terpinene (~18%), and para cymene (~11%) were detected as the most abundant molecules. Other compounds such as alpha pinene, alpha thujene, sabinene, beta pinene, alpha terpinene, limonene, 4-thujenol, o-isopropenyl toluene, caryophyllene, bicyclo (5.3.0) decane, 2- methylene, alpha guaiene, alpha panasinsen and caryophyllene oxide were also present in relatively small amounts.

**Figure 1.**
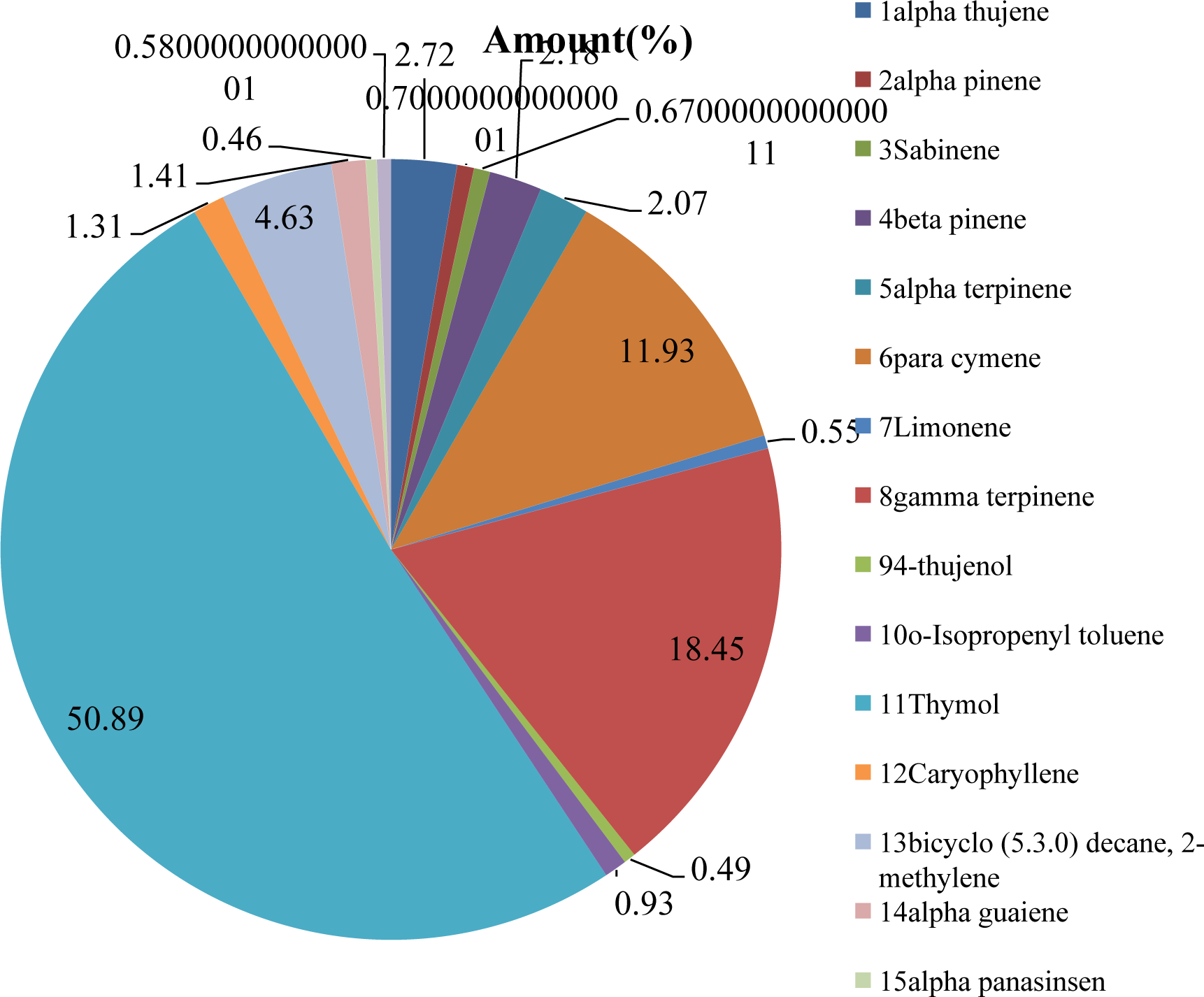
Chemical profile of leaf essential oil of O.viride. The GC-MS ananlysis of the EO showed the presence of thymol as the major constituent.

### Antibacterial activity

The antibacterial activity assay of the EO against five gram positive and five gram negative bacterial strains and obtained the results as shown in Table 1 **(Table 1)**. The EO strongly inhibited the growth of *Micrococcus luteus* (zone of inhibition; 11±0.4mm)*, Bacillus subtilis* (zone of inhibition; 14±0.5mm)*, Psudomonas alcaligenes* (zone of inhibition; 10±0.6mm), *Bacillus cereus* (zone of inhibition; 12±0.6mm); while moderate growth inhibitory activity was observed for *Staphylococcus aureus* (zone of inhibition; 04±0.2mm), *Enterococcus faecalis* (zone of inhibition; 06±0.2mm)*, Pseudomonas aeruginosa* (zone of inhibition; 05±0.3mm), *Alcaligenes denitrificans* (zone of inhibition; 06±0.3mm), whereas, *Escherichia coli* and *Campylobacter coli* have not exhibited significant sensitivity to essential oil. Chloramphenicol was used as reference antibiotic against all test organisms. Our results revealed that EO possesses potent antimicrobial activity against the broad range of gram positive and gram negative bacteria. These results are in agreement with the already existing literature citing the antimicrobial activity of essential oil of *O. viride* and *Ocimum gratissimum* against *Listeria monocytogenes* [11].

**Table 1.** Antibacterial activity of leaf essential oil of *O.viride*.

| Antibacterial assay |  |  |
| --- | --- | --- |
| Zones of inhibitions (mm) |  |  |
| Bacterial strains | EO (0.1µl/ml) | Cmp+ |
| 1. *Micrococcus luteus* | 11±0.4 | 25±1.05 |
| 2. *Staphylococcus aureus* | 04±0.2 | 18±0.5 |
| 3. *Escherichia coli* | - | 15±0.5 |
| 4. *Enterococcus faecalis* | 06±0.2 | 18±0.43 |
| 5. *Campylobacter coli* | - | 28±0.9 |
| 6. *Bacillus subtilis* | 14±0.5 | 15±0.55 |
| 7. *Bacillus cereus* | 12±0.6 | 08±0.25 |
| 8. *Alcaligenes denitrificans* | 06±0.3 | 21±0.75 |
| 9. *Pseudomonas aeruginosa* | 05±0.3 | 16±0.09 |
| 10. *Pseudomonas alcaligenes* | 10±0.5 | 15±0.6 |

### Molecular docking analysis

To understand the possible mechanism behind the antibacterial activity of EO against the bacteria *Bacillus subtilis*, we performed the molecular docking analysis of the major constituent in the essential oil, thymol against various bacterial proteins. It was found that thymol shows the best binding affinity with bacterial secA protein. Where, secA protein is an ATPase that drives the post-translational translocation of proteins through the secY channel in the bacterial inner membrane and possibly acts possible receptor for the ligand thymol. *In silico* molecular docking analysis was carried out to find an orientation of lead molecule in target protein. Although molecular docking screens large databases of small molecules by orienting and scoring those in the binding site of a protein. Top-ranked molecules may be tested for binding affinity *in vitro* and may become lead compounds, the starting point for drug development and optimization. With the protein secA ATPase, an interaction was observed with minimal H-Bond energy of −2.5kcal/mol by many components of different analogs of thymol followed by the scoring function in order to evaluate the interactions between sec A ATPase and thymol. Docking energy was used to identify the correct binding pose and then ranked the most befitting target ligand complex based on their binding affinity and RMSD value. In this work docking was performed with the help of software Molegro Virtual Docker (MVD) for predicting the binding energy of thymol with interaction between secA protein. Different analogs of thymol were used but the best analog (zinc id-967515) on the basis predicted binding energy and other binding parameters like hydrogen bond interaction and electrostatic interaction was selected. After docking thymol was found to interact with protein sec A with binding energy **-**15 kcal/mol with the active site Lys284, Trp275, Leu269, Arg19,Glu277, pro270 **(Fig 2)**. Thymol can serve as a ligand for the drug target and in the future and can be optimized to form an improved antimicrobial drug.

**Figure 2.**
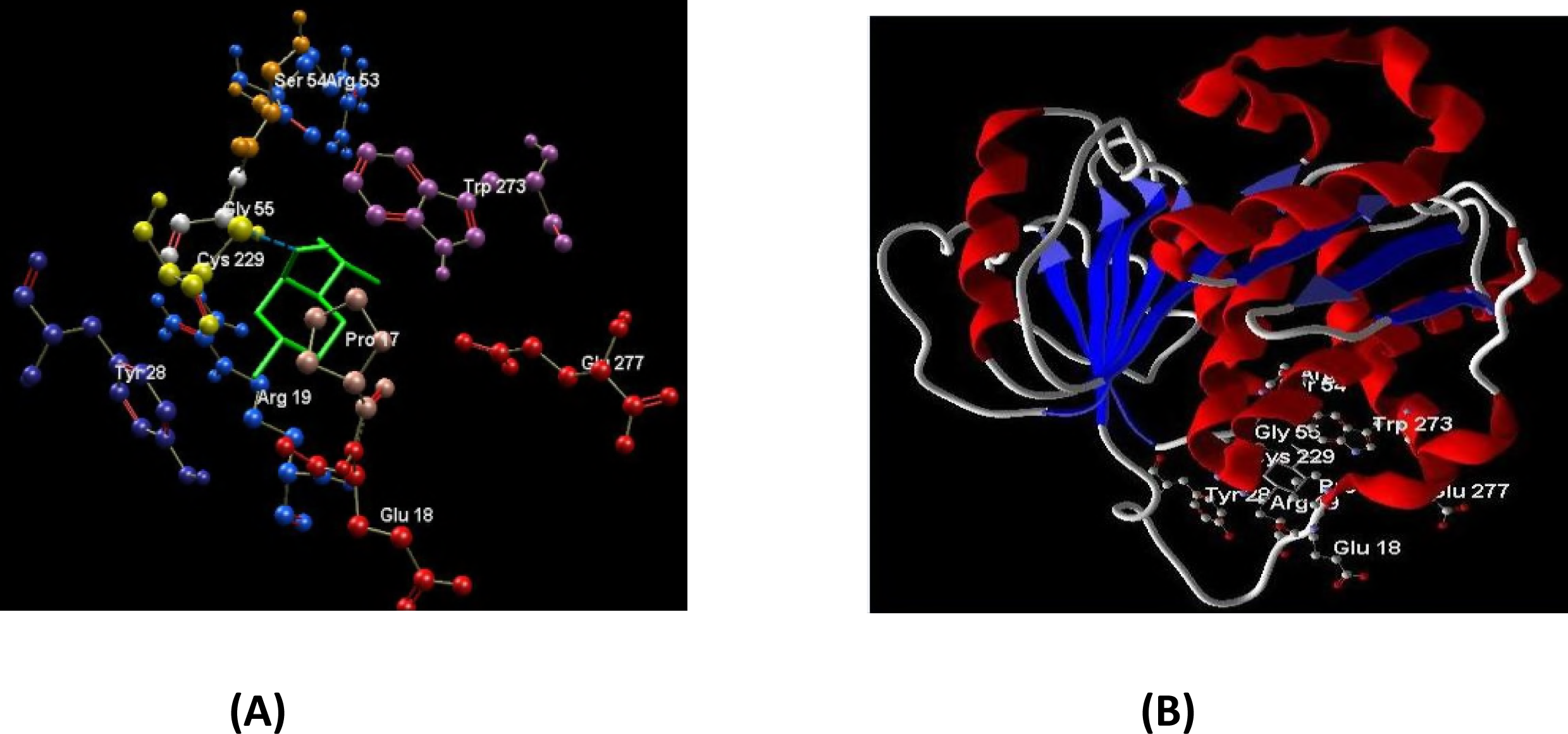
Molecular docking analysis of thymol against the bacterial Sec A ATPase. **(A)** Green ligand is surrounded by amino acid residues with which ligand has hydrogen bonds. The amino acids are labeled as xxx123. In the binding moiety Lys, Trp, Leu, Arg, Glu and Pro are placed at 284, 275, 269, 19, 277 and 270 position respectively.**(B)** Docking view of 2-Isopropyl-5-methylcyclohexanol against secondary structure of 3RPH with red colour showing helix.

### *In vitro* cytotoxic activity

Colorimetric MTT assay was done to elucidate the cytotoxic potential of the leaf essential oil of *O. viride* with different concentrations (0.01-0.1μl / ml). A dose dependent inhibition by essential oil was observed with IC_50_ values of 0.042±0.004μl/ml, 0.052±0.006μl/ml, 0.045±0.002μl/ml, 0.034±0.001μl/ml, 0.040±0.005μl/ml and 0.064±0.007μl/ml against prostate DU-145, liver HEP-2, neuroblastoma IMR-32 and colon HT-29, 502713, SW-620 respectively **(Fig 3)**. DMSO (0.02%, v/v), did not affect the cell growth when treated for the same time period. The cytotoxicity of essential oil was also determined using normal monkey kidney cell line (CV-1) and no significant cytotoxicity was observed (IC_50_ - 1.92μl/ml). The apoptotic activity of the essential oil isolated from several Lamiaceae members have been reported in the literature [8,10]. The observed cytotoxicity might be due to the presence of various chemical components in the essential oil including monoterpenes such as thymol. Monoterpenes have been reported to exert antitumor activities and suggest that these components are a good source of cancer chemopreventive agents [20].

**Figure 3.**
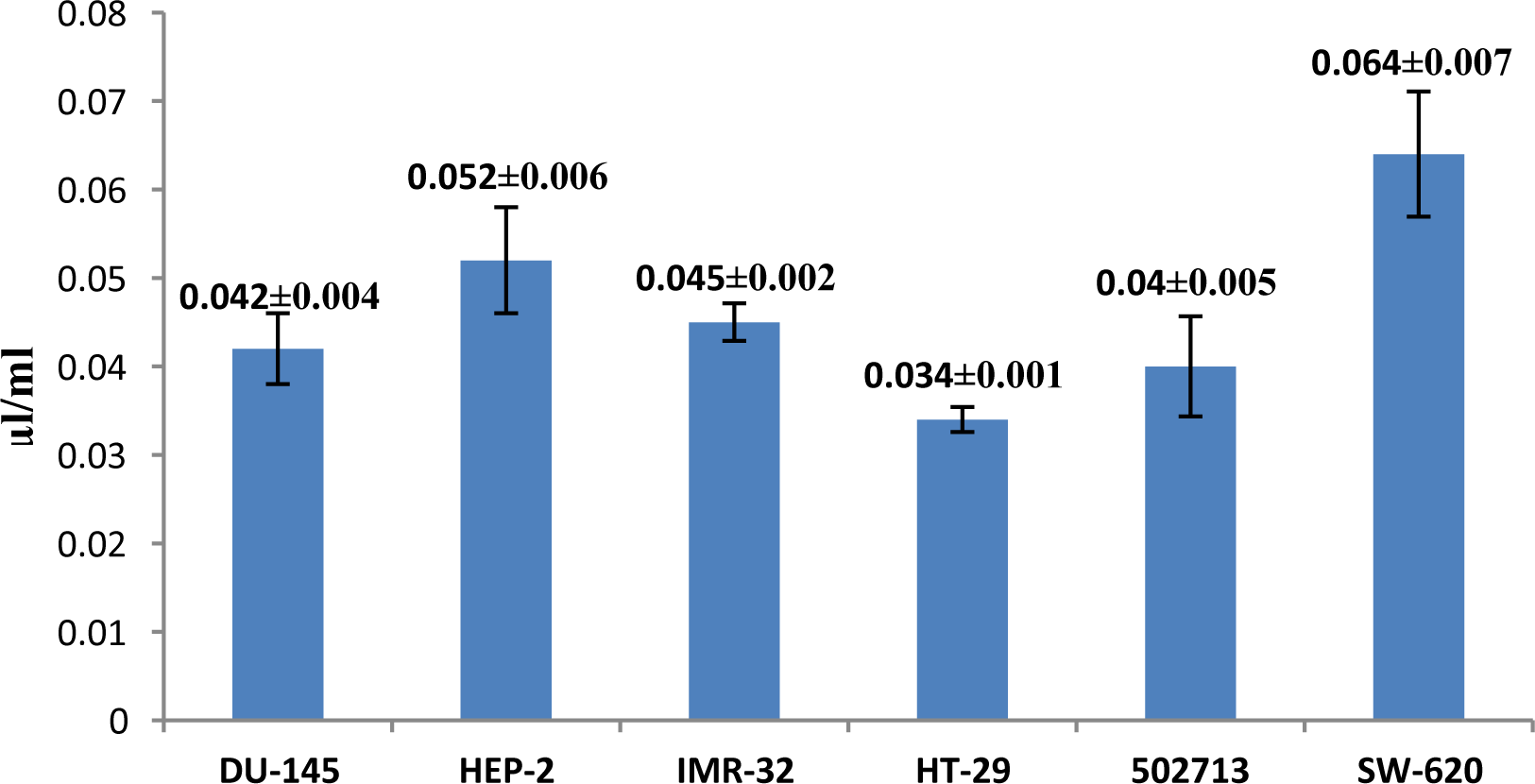
Cytotoxityactivity of the leaf essential oil of O.viride. The EO was found to be highly effective against the HT-29 colon cell line with respect to other prostate DU-145, liver HEP-2, neuroblastoma IMR-32 and colon 502713, SW-620.

### Cell cycle analysis

The cell cycle progression to different concentrations of essential oil was studied, by using PI staining method. The untreated colon HT-29 cells showed 7.69% of cells in the sub G1 phase **(Fig 4a)**. When treated with the essential oil of *O. viride* (0.1μl/ml), the HT-29 cell population in the sub G1 phase increased significantly up to 78.16% **(Fig 4b)** thereby showing more cell death in the sub G1 phase. Drug camptothecin (1μM) was used as control to further validate our experiment **(Fig 4c)**.

**Fig 4.**
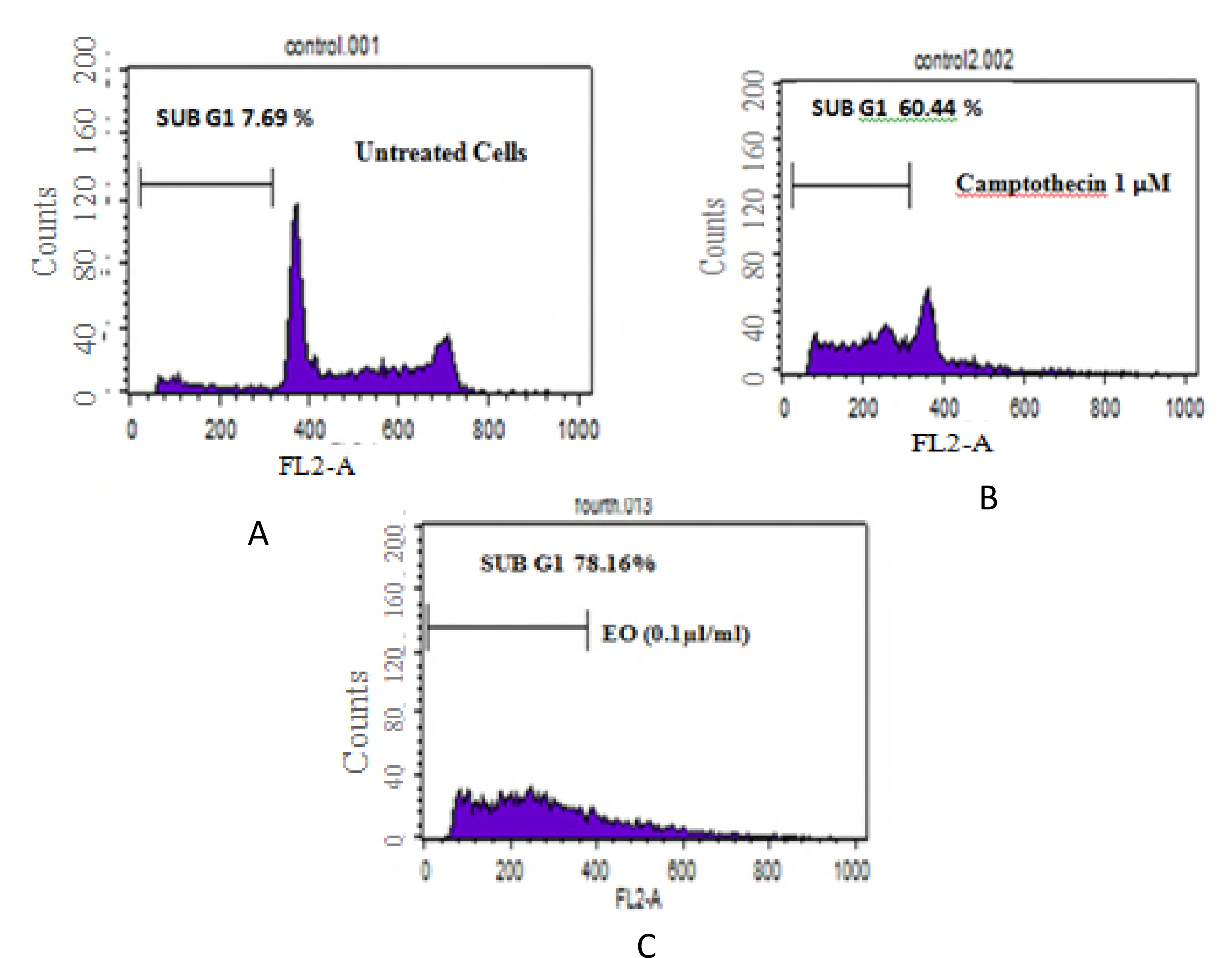
Cell cycle analysis of HT-29 cells. (**A**) Untreated HT-29 cellsshowing 7.69% cells in the SUB G1 phase. (**B**)Positive control-cells treated with camptothecin (1μM) showing 78.16% cells in the SUB G1 phase. (C) HT-29 cells treated with 0.1μl/ml essential oil showing 78.16% cells in the SUB G1 phase.

### Effect of essential oil on determination of morphological changes in HT 29 cells

Nucleic acid staining with Hoechst 33258 revealed characteristic apoptotic nuclei, that exhibited highly fluorescent condensed chromatin in cells treated with the essential oil (**Fig 5**). The morphological changes and cell death of HT 29 cells were observed at concentrations of 0.1μl/mL. Most cells were detached from the dishes, and cell rounding and shrinking occurred at the same concentration of the essential oil (**Fig 5**).

**Fig 5.**
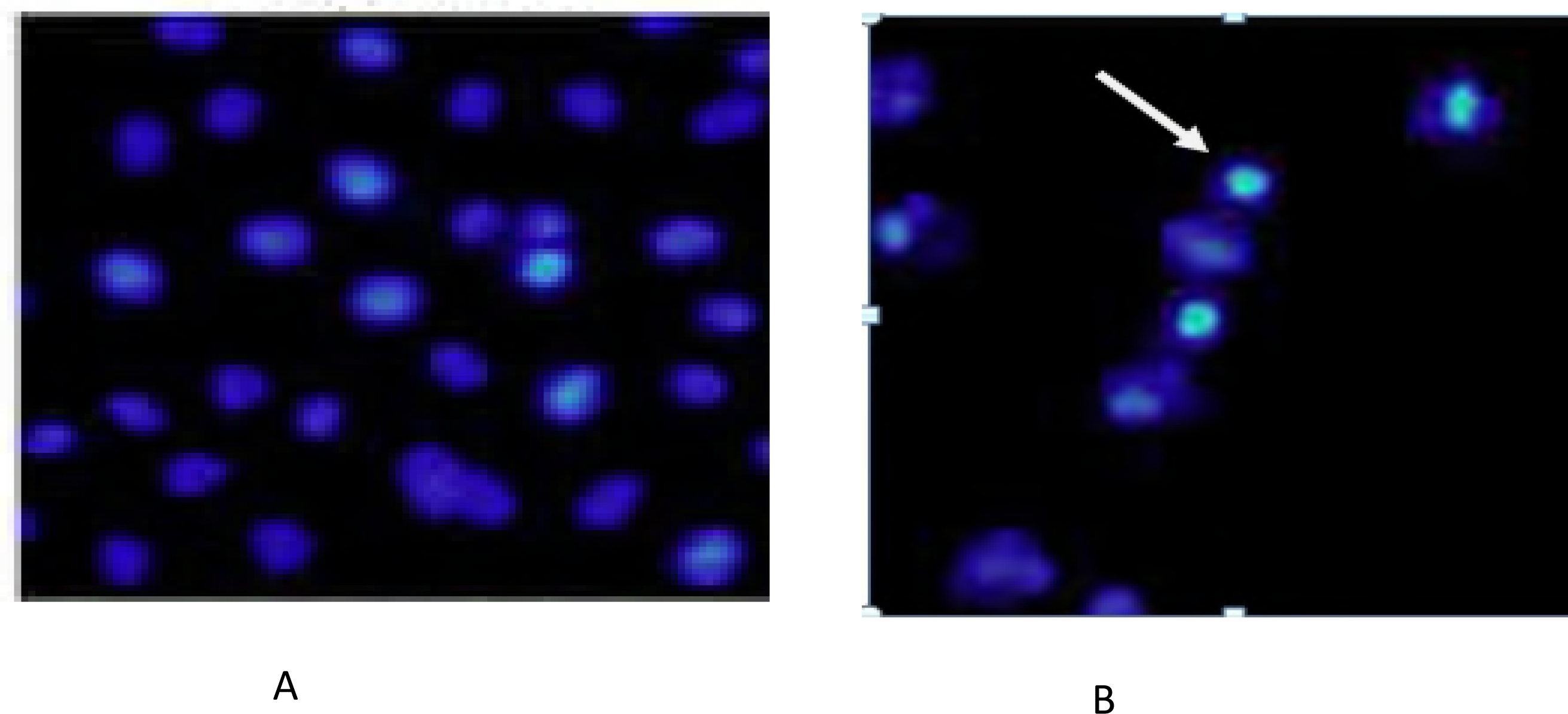
DNA fragmentation assay in colon HT-29 cells. Lane 1 (Untreated cells), Lane (Camptothecin 1 μM), Lane 3 (Camptothecin 1 μM), Lane 4 (*O.viride* essential oil 0.1 μl/ml).

### DNA fragmentation assay

To understand whether the essential oil would induce apoptosis in HT 29 cell line we assessed the DNA fragmentation induction, a biochemical hallmark marker for the induction of apoptosis *in vitro*, we observed essential oil induced endonucleolytic DNA cleavage in a dose-dependent manner in genomic DNA isolated from colon HT-29 cells and efficient induction of apoptosis was observed at 0.1μl/ml of essential oil **(Fig 6)**.

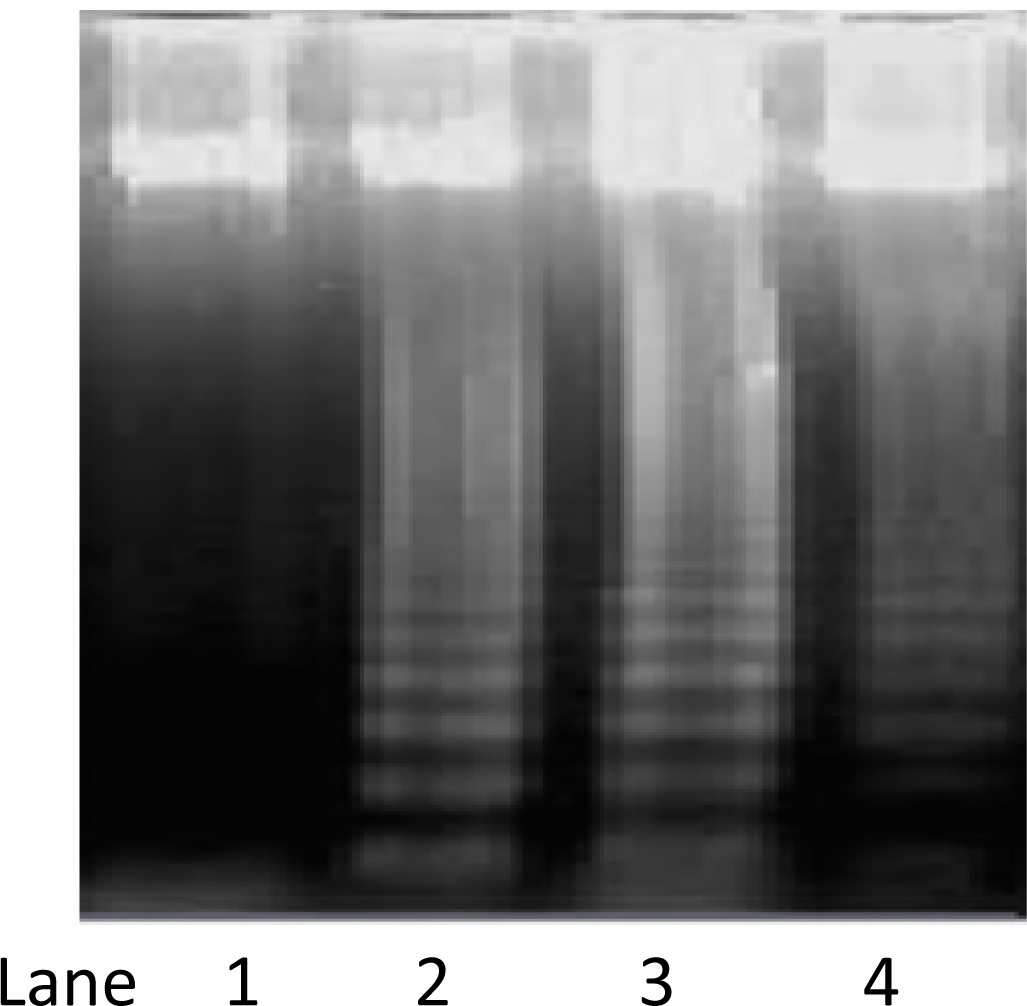

## Discussion

Development of effective therapeutic agents against cancer and infectious diseases remains main concern till today and approaches are more focused towards the natural product drug discovery. Plant products are well known for the treatment of cancer and other diseases, among various categories essential oils recorded as complex mixture of secondary metabolites like monterpenes and sesquiterpenes and are reported to induce sensitive growth inhibition against microbes as well as against cancer cells [7]. Induction of apoptosis is a highly desirable goal for cancer control. Apoptosis regulators have been suitable targets for cancer therapy over several decades. The significant toxicity against microbes and cancer cell shown in our study by *Ocimum viride* may be due to the presence of the major compound thymol, a terpene, has been extensively explored as a potential antibacterial agent against many bacterial strains [26, 27, 28]. In addition, large number of other aromatic components mainly thymol, eugenol, cymene limonene etc., are also reported by other researchers [15] as well as from our studies from the essential oil of *Ocimum viride,* it is also possible that the component in lower percentage may also be involved in some type of synergism with other active or mojor compounds. Our analysis showed the highest sensitivity of the essential oil towards *Bacillus subtilis* pathogen (14±0.5mm) and to correlate the relation between the thymol content and its antibacterial activity as earlier reported by various research groups, *in silico* approach by molecular docking was carried out involving various bacterial proteins of *B. subtilis* that were docked with thymol to predict the best binding efficiency. The molecular docking of thymol with bacterial sec A protein resulted in best binding energy of −15 kcal/mol with the active site Lys284, Trp275, Leu269, Arg19, Glu277 and pro270. Since, bacterial secA is an ATPase that is responsible for post translational translocation of proteins and possibly serving as receptor for thymol thereby causing interference in protein transport in bacteria [27]. Further, our investigation lead us to the understanding that *Ocimum viride* essential oil was also found effective in combating the cell proliferations through induction of apoptosis of HT-29 colon cancer cells. Apoptosis as understood is recognized as choice of physiological cell death and is accompanied by a specialized series of cellular events such as chromatin condensation, DNA fragmentation, cytoplasmic membrane blebbing and cell shrinkage [10, 26, 30].

The specific action of the essential oil toward cancer cells might be partially related even to cell mitotic rate. Indeed, HT-29 colon cell line was found most sensitive to essential oil administration among others, also presented the lowest doubling time (approximately 24 hours) among the cell lines tested, possibly indicating that some EO components (such as thymol) might interfere with cell cycle and DNA synthesis. The sub-G1 peak, of cell cycle analysis bears cell population containing apoptotic nuclear fragments such as apoptotic bodies, chromatin condensation and DNA fragmentation into oligonucleosomal-sized fragments that give indication of the regulated cell death. The similar findings were also reported on COLO 205 colon cancer cells [32]. Although diverse phytoconstituents have been reported from various species of *Ocimum* for various biological processes including anticancer potential [33,34,35,36,37] but it cannot be ruled out that the antimicrobial and anticancer activity shown by the essential oil may be due to the synergistic or antagonistic effects of identified monoterpene compounds and sesquiterpene as reported earlier [38]. Further there is still need to understand the detailed mechanism of action of essential oil against cell lines at both cellular and molecular level which can lead in the direction of more efficient and sustainable release of essential oil.

## Conclusions

*Ocimum viride* essential oil was isolated from aerial part that were chemically analyzed and evaluated for antimicrobial and antiproliferative efficacy. Essential oil demonstrates antiproliferative efficacy against HT-29 colon cancer cell line via induction of apoptosis and potential antibacterial activity against *Bacillus subtilis* was also established by our studies. The major compound identified as Thymol, gamma-terpinene, para cymene etc. Thymol was found to have appropriate binding energy with the bacterial sec A ATPase protein indicating the involvement of the membrane disruption for antibacterial activity. Overall, the data showed the importance of preliminary bioassays as a screening of the bioactive from plant products and in this study an essential oil from *Ocimum viride* established its importance as potential source of bioactive compounds.

## Acknowledgements

Authors would like to acknowledge the School of Biotechnology and Department of Bio-informatics, University of Jammu, for their support. Here, we would also like to mention the work is possible due to the facility created by the funding of DST-FIST, UGC-SAP, RUSA, PURSE. The authors also wish to acknowledge IIIM-CSIR, Instrumentation division and Cancer pharmacology division Jammu for extending their facilties.

## Authors Contributions

Conceptualization/ Supervision/ Methodology/ Writing: MB Writing/Review writing: MS Software analysis: KA Editing: RA

